# Behavioral and causal evidence for object-based scene recognition in visual cortex

**DOI:** 10.64898/2026.06.29.735096

**Authors:** Aaron Schnippe, Natalia Rutkowska, Marius V. Peelen, Marco Gandolfo

## Abstract

Our visual environment can be parsed into objects and scenes, a distinction that is reflected in the organization of the human visual cortex. Previous research has shown that object and scene perception nevertheless closely interact, such that scenes influence object perception and objects influence scene perception. It remains unclear, however, whether and how objects that are not inherently diagnostic of their surroundings aid the recognition of poorly visible scenes (e.g., a person standing in a dark living room). Here, in three behavioral experiments, we show that participants made more accurate indoor/outdoor judgments when degraded scene photographs were presented together with an object than when the scene or the object was shown alone, even though the same object categories appeared in indoor and outdoor scenes. This object-driven benefit vanished once scene structure was removed through phase scrambling and was reduced when objects appeared in physically inconsistent locations within the scenes. These results suggest that objects in consistent locations (e.g., a person standing on a floor) disambiguate scene layout. Finally, in a pre-registered transcranial magnetic stimulation (TMS) study (N = 48), we provide causal evidence that the object-selective lateral occipital cortex (LOC) supports scene categorization when scene layout is disambiguated by within-scene objects. Stimulation of the LOC, particularly at 260-300 ms after stimulus onset, selectively disrupted object-based scene recognition. Together, these findings demonstrate that objects facilitate the read-out of the surrounding space in service of efficient scene recognition.

**Significance Statement:** Understanding how scene and object processing interact for efficient recognition is a key question in natural vision. Research has long emphasized how surrounding scenes help us identify objects, yet the reverse – how objects shape the perception of scenes – has received little attention. In the dark, does a glimpse of a floating boat tell us we are looking at a lake? In this study we demonstrate that a single object helps people recognize hardly visible scenes. This benefit required intact scene structure and depended on where the object appeared in the scene. In addition, object selective visual cortex was causally related to this benefit. Together, these findings show that objects’ visual appearance can be used to better understand our surroundings.

## Introduction

The visual environment consists of scenes and objects. Although their visual processing is separable in the brain (Grill-Spector, 2003; Park et al., 2011; Dilks et al., 2013; Epstein & Baker, 2019; Wischnewski & Peelen, 2021A, 2021B), scenes and objects are not processed independently of one another.

Much of what is known about these interactions concerns how scenes influence object processing. Objects are easier to recognize within a semantically congruent scene (e.g., a car on a road versus a car on a beach; Biederman et al., 1982; Boyce et al., 1989; Davenport & Potter, 2004; Davenport, 2007). Heavily blurred objects are likewise recognized more accurately, and even appear sharper, when embedded in a scene rather than shown in isolation (Brandman & Peelen, 2017; Rossel et al., 2022), underscoring the importance of scene context for object recognition.

In the brain, scene context enhances the multivariate representation of poorly visible objects in the object-selective cortex (OSC), including the lateral occipital cortex (LOC) (Brandman & Peelen, 2017), an effect that peaks around 320 ms after stimulus onset (Brandman & Peelen, 2017; Leticevscaia et al., 2024). A chronometric transcranial magnetic stimulation (TMS) study further showed that both object-and scene-selective regions causally contribute to recognition when an object’s identity is disambiguated by the surrounding scene. Together, these findings suggest that interactions between the scene-selective cortex (SSC) and OSC drive behavioral scene-context effects, highlighting their perceptual rather than post-perceptual nature (Bar, 2004; Peelen et al., 2024).

The reverse interaction — how objects influence scene recognition — has received far less attention. Object-scene relationships can either improve or impair scene recognition, depending on congruence (Davenport & Potter, 2004; Davenport, 2007; Joubert et al., 2007; Leroy et al., 2020; Furtak et al., 2022). Indeed, when an object is sufficiently diagnostic (e.g., a bath), even a scene (e.g., a bathroom) reduced to that single object can be categorized with high precision (Wiesmann & Võ, 2022).

Objects may also help disambiguate scenes that are hard to recognize irrespectively of the semantic relationship between the object and the scene. A cat, for instance, sitting in a dark room may define the perspective of a scene, facilitating to locate where the floor and the walls are (See **Figure 1**). Consistent with this, an fMRI study found that classifiers trained to distinguish intact scene categories (indoor vs. outdoor) in the SSC generalized better to blurry scene layouts containing a disambiguating object than to layouts shown without their object, or to the object alone on a uniform grey background (Brandman & Peelen, 2019). A subsequent MEG study using the same stimuli showed that the object’s influence on scene representations peaked 320 ms after stimulus onset (Brandman & Peelen, 2023). These results suggest that objects can disambiguate a scene’s layout, and that this process is reflected in visual cortex responses.

**Figure 1.**
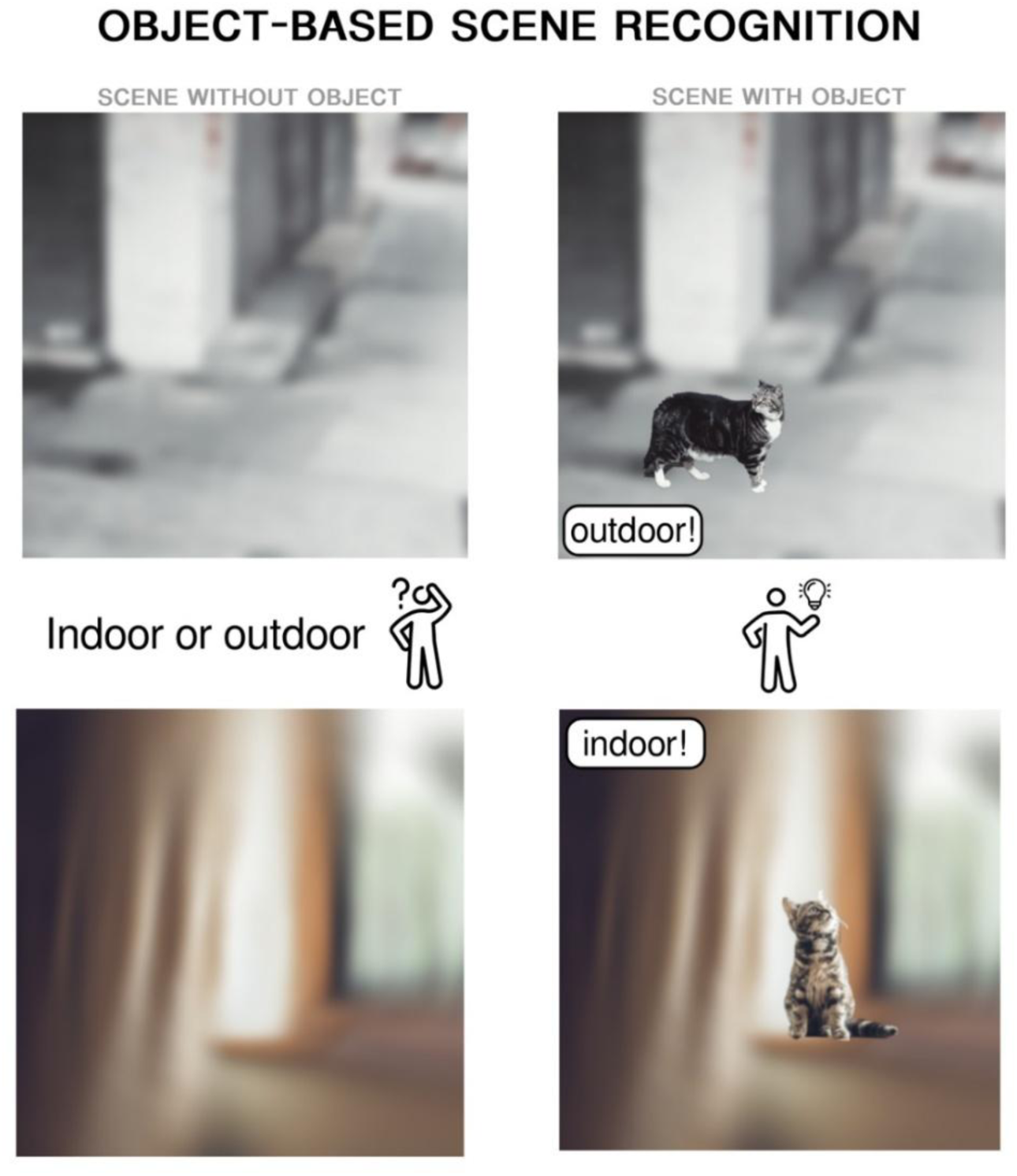
**Illustration of object-based scene recognition**. On the left, degraded photographs of scenes with their object (originally belonging to the photographs) removed. These scenes are hard to categorize as indoor or outdoor. On the right, the same scenes with their object. The object is not in itself diagnostic of the scene category (cats occur both indoors and outdoors). In the current study, we hypothesize that the object can improve scene recognition via a disambiguation process. Disambiguation of the poorly visible scene can occur for example by the object specifying the location and orientation of the floor surface and adding depth information and perspective, ultimately facilitating the recognition of the environment as indoor or outdoor.

However, the object-based scene recognition observed in visual cortex through these correlational methods may be epiphenomenal, potentially unrelated to how people actually arrive at correct scene judgements. A recent behavioral study suggested that object information may sharpen the appearance of the surrounding scene, as measured by a blur-matching paradigm (Carrez-Corral et al., 2025). Yet it remains unclear whether this object-based impression of increased visibility translates into improved scene recognition, and whether such effects relate directly and causally to visual cortex activity.

To address these questions, we ran four experiments testing whether objects support the recognition of degraded scenes. In Experiment 1, we compared scene-categorization judgments (indoor vs. outdoor) for ambiguous scenes presented with their object, without their object, or as the object alone; objects improved the recognition of ambiguous scenes. In Experiments 2 and 3, we show that this benefit is not driven by low-level visual features of the ambiguous scenes and that it is sensitive to the object’s original position within the scene. Finally, in Experiment 4, using TMS, we show that LOC is causally related to the behavioural effects of object-based scene recognition.

## MATERIALS AND METHODS

All data, stimuli and analysis scripts necessary to reproduce the reported results are available via the Open Science Framework: https://osf.io/teuxv/overview?view_only=d46fb6f0a8784dcf9e19c6880bf228c9. For Experiment 4, pre-registration is available via aspredicted.org (https://aspredicted.org/m9gp-k5hj.pdf).

### Participants

Experiment 1-3 were conducted online. Participants were recruited via Prolific (Prolific Inc.), received monetary compensation (∼£8/h) and the procedures were approved by the Radboud University Ethics Committee (ECSW – 2022-079). Experiment 4 was conducted in the laboratory, and participants were recruited via the university’s participant panel (SONA systems). The study procedures of Experiment 4 were approved by the’Centrale Commissie voor Mensgebonden Onderzoek (CCMO)’ under project number 2020-6140 (NL72752.091.20) and conducted in accordance with the Declaration of Helsinki. Participants were monetarily compensated (15 euros/h) unless they specifically requested for course credits.

### EXPERIMENT 1

We planned a sample of 50 participants. A sensitivity analysis showed that this sample is sufficient to detect a small to medium effect size in a one-way repeated measures ANOVA with three levels (partial η^2^ = 0.03) with a power of β =.80 and a significance level of α = 0.05. In total, we tested sixty-one participants (33 females, 28 males; mean age 36.27 years; *SD* = 13.24). Eleven participants were excluded during the analysis due to performance at chance level (see Analysis section), so that the final sample size consisted of the 50 planned participants (27 females, 23 males; mean age 35.17 years; *SD* = 12.64).

### EXPERIMENT 2

We planned a sample of 50 participants, similar to Experiment 1. Fifty-eight participants participated in the experiment (32 females, 26 males; mean age 36.45 years; *SD* = 13.76). After removing participants performing at chance, the final sample consisted of 51 participants (29 females, 22 males; mean age 37.10 years; *SD* = 14.32).

### EXPERIMENT 3

Because one extra factor was added to the design (object position), and because we expected that the effect of object position on scene recognition would likely be smaller, we aimed for a sample size of n = 100. A sensitivity analysis on a paired two-tail contrast expressing the interaction effect revealed that a sample size of n = 100 is sufficient to detect an effect of d = 0.25 with a power of β =.80 and a significance level of α = 0.05. One-hundred and four participants took part in the experiment (32 females, 70 males, 1 identified as other, 1 preferred not to say; mean age 36.74 years; *SD* = 11.27). Four participants were excluded during the analysis due to performance at chance level so that the final sample consisted of 100 participants (31 females, 67 males, 1 identified as other, 1 preferred not to say; mean age 36.89 years; *SD* = 11.25).

### EXPERIMENT 4

We tested a total of 53 participants (36 female, 17 male, mean age 22.91 years; *SD* = 3.48). After exclusions, our final sample included 49 participants after exclusions (See analyses; 32 females, 17 males; mean age 22.96 years; *SD* = 3.54 years). We preregistered a sample size of 48 healthy participants, 24 per stimulation site. This sample size matches the one used by Wischnewski and Peelen (2021A), a TMS study investigating context-based object recognition using a comparable design. One additional participant was tested because they had already completed the localization session (see below). Adding this participant to the analyses did not change the overall pattern of results. All participants had normal or corrected-to-normal vision and were right-handed. Participants with neurological or psychiatric disorders, CNS-acting medication, a (family) history of epilepsy or convulsions or seizures, brain surgery, cochlear or metallic implants in their head or neck area, cardiac pacemaker or intra-cardiac lines, a medication infusion device, a neuro-stimulator, or pregnancy were screened-out from participation.

### Stimuli

#### EXPERIMENT 1

The experiment included 96 unique scenes, 48 photographs of outdoor (e.g., beaches, gardens, forests) and 48 photographs of indoor (e.g., kitchens, living rooms, malls) environments with one main object in the foreground. These objects belonged to one of six categories (cat, dog, human, chair, lamp, plant). Objects of these categories appeared in both indoor and outdoor scenes, so that object category was not diagnostic of scene category. The object was temporarily cropped out and the scenes were degraded using Gaussian and radial blur. The scenes were additionally desaturated to reduce the effect that colour alone can have on scene categorization (Oliva & Schyns, 2000; Yao & Einhäuser, 2008). Each image was exported in three ways (see **Figure 2A**): the degraded scene including the object (“scene-with-object” condition), the degraded scene without the object (“scene-without-object” condition), and the object on a uniform grey background (“isolated-object” condition). Thus, the final stimulus set contained 288 images across all conditions.

**Figure 2.**
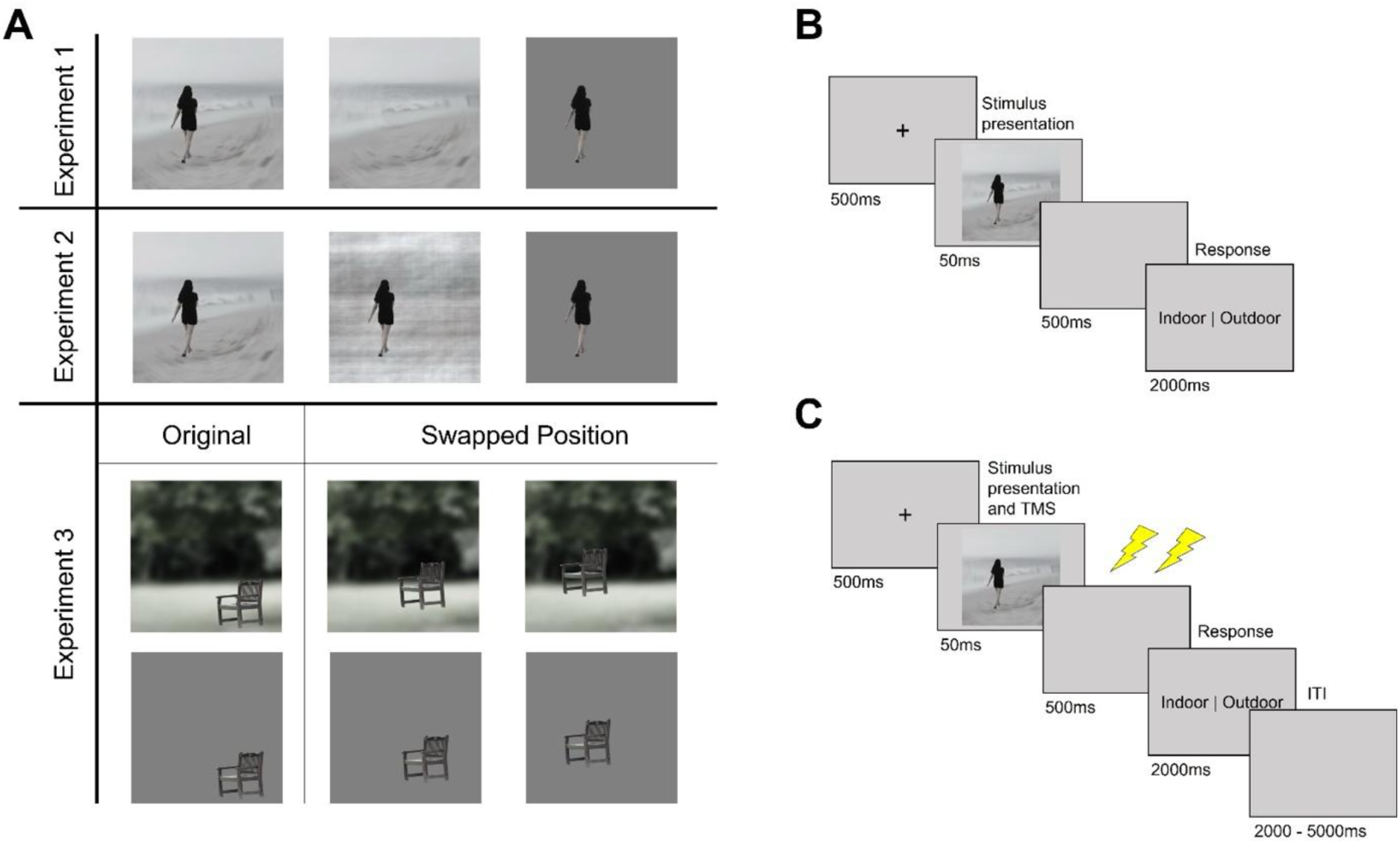
(A) Example stimuli used in the experiments. Stimuli could show indoor or outdoor scenes. The objects within the scenes belonged to one of six categories (cat, dog, human, chair, lamp, plant). These categories appeared in both indoor and outdoor scenes, so that object category was not diagnostic of scene category. Each stimulus was created in three conditions (left to right): A degraded scene including the object; a degraded scene without the object; the object on a uniform background. In Experiment 2, object-and scene-categories were identical to Exp 1, but the isolated scene condition was replaced with a condition depicting the object in a phase-scrambled version of the scene. In Experiment 3 the object in each stimulus was either shown at its original position in the scene, or at the position that another object has in its respective scene (“swapped position”). (B) Procedure of Experiments 1-3. After a fixation cross (500ms), the stimulus was presented for 50ms. Then a blank screen appeared (500ms), followed by a 2000ms response window (the indoor/outdoor labels are presented for illustrative purposes, but were not presented in the actual experiment). The participant responded using the “f” or “j” key on the keyboard. (C) Schematic trial for Experiment 4. After stimulus onset, participants received double pulse TMS (25 Hz) over LOC or OPA at one of three time points (60ms after stimulus onset, 160ms after stimulus onset, 260ms after stimulus onset).

#### EXPERIMENT 2

A subset of 66 scenes of Experiment 1 (33 indoor, 33 outdoor photographs) was used. In this experiment we generated a phase-scrambled version of each scene (without object) and then placed the object on top of this background (“scrambled-scene-with-object” condition – **Figure 2A**). In this condition, the object was temporarily cropped out, and the scene was then degraded through phase scrambling (Thomson, 1999) using a Matlab function (https://martin-hebart.de/webpages/code/stimuli.html). The final images (198 in total) included a degraded scene with an object, the object on a neutral background and the phase-scrambled scene with an object.

#### EXPERIMENT 3

We used a subset of 64 scenes (32 indoor, 32 outdoor photographs). Starting from these, we then generated images in which the object position was swapped to the position of any of the other objects in our image set (“swapped position,” see **Figure 2A**). For example, if object A had position coordinates xA/yA in its original Scene A, and Object B had position coordinates xB/yB in its original Scene B. In the swapped position condition, Object A was presented in Scene A at position coordinates xB/yB. This was done to keep the average position of the objects the same within the original and swapped stimulus set, and to ensure that the object would still appear in a different, but plausible position in the scenes. For each participant, the swapped stimulus set was created by randomly assigning to an object the position of the object of another scene.

#### EXPERIMENT 4

The same 64 indoor and outdoor scene photographs as in Experiment 3 were used. The scene-with-object condition showed the object in its original position on the degraded scene – exactly like in Experiments 1-3. Differently from the other experiments, the scenes without objects here were made sharper and less degraded with the aim to match, on average, the behavioral performance of indoor/outdoor judgments between the two conditions. This was done to compare TMS effects across conditions within the same range of performance, avoiding ceiling and floor effects. To achieve that, we ran a validation pilot experiment online (N = 29), contrasting performance on scene judgments for the scene-with-object vs the scene-without-object in its less blurry version using the exact same procedure as the TMS experiment (i.e., same intertrial interval; see below) and found no significant differences between these two conditions (*t*(28) = 1.654, *p* =.109). Stimuli were displayed on a BenQ Mobiuz 27” 120Hz computer screen with a fixed size of 10×10 cm.

### Transcranial Magnetic Stimulation

#### EXPERIMENT 4

TMS was applied via a C-B60 figure-of-8 coil with an outer diameter of 75 mm, which received input from a Magpro-X-100 magnetic stimulator (MagVenture, Farum, Denmark). In two participants, a MC-B70 butterfly-shaped figure-of-8 coil with an outer diameter of 96 mm was used instead. Participants received double-pulse TMS at a stimulation intensity adjusted to 85% of the individual phosphene threshold of each participant. The double-pulse was given at 25 Hz (two pulses 40 ms apart). The phosphene threshold was determined by increasing stimulation intensity during single-pulse stimulation of the early visual cortex (defined by placing the stimulator 2cm above the inion) until visual phosphenes were reported in 50% of the trials while participants fixated on a grey screen in a dimly lit room. To further ensure that participants were seeing phosphenes, we also shifted the stimulator position to ensure that we were able to elicit phosphenes in the contralateral hemifield. The average threshold value was 56.4% of the maximum stimulator output. The TMS coil was placed with the help of an infrared-based neuronavigation system (Localite, Bonn, Germany) using an individually adapted standard brain model over the left lateral occipital cortex (LOC), selective for objects, or left occipital place area (OPA), selective for scenes. The stimulation site in each participant was chosen based on a localization session that took place 3 to 7 days before the experiment, following Wischnewski and Peelen (2021A, 2021B, see Supplementary Materials for details). The stimulation coordinates for LOC and OPA were identified through Talairach coordinates set in the Localite neuronavigation system. The coordinates used for the left LOC were-45,-74, 0 and for the left OPA, the coordinates were-34,-77, 21. Note that the coordinates used were the left-hemispheric homologue coordinates of the LOC (Pitcher et al., 2009) and OPA (Julian et al., 2016) defined in the right hemisphere. The stimulation of the left hemisphere is based on the results of Brandman and Peelen (2019), who found neural evidence for object-based scene recognition in the left hemisphere. TMS pulse delivery relative to the onsets was controlled via Matlab through a USB to TTL port connected to the stimulator with the help of functions of the MAGIC toolbox (Habibollahi Saatlou et al., 2018).

### Procedures and design

#### EXPERIMENT 1 - 3

On each trial, participants were presented with a fixation cross (500ms), followed by the stimulus presentation (50ms), a blank screen (500ms), and a two-second two-alternative forced choice response window (**Figure 2B**). Before the recorded trials, participants were given instructions and completed a practice block of 12 trials (4 trials for each of the three conditions) using stimuli that were not used in the main experimental block. During the practice, participants received feedback after each trial (“correct” or “incorrect”). Participants did not receive direct feedback during the experiment. However, every 24 trials, a break screen appeared presenting the average response time and accuracy of their last 24 trials. For all participants, the stimuli were fit to a size of 10×10 cm, irrespective of the monitor’s resolution.

Experiment 1 included three within-subject conditions: scene-with-object, scene-without-object, and isolated object (**Figure 2A**). Each participant completed two blocks of 144 trials presented in random order. To avoid familiarity and carry over effects, the trial list across participants was split to present half of the scenes in the scene-with-object condition (48 scenes) and the other half in the isolated object and scene-without-object conditions (i.e., the other 48 scenes in these two conditions). The assignment was counterbalanced across participants.

In Experiment 2, instead of the scene-without-object condition, participants were presented with the scrambled-scene-with-object condition. Further, participants were never presented with the same image in different conditions. This resulted in three sets of stimuli, each consisting of 22 stimuli in the scene-with-object, scrambled-scene-with-object and isolated object condition, and thus 66 stimuli in total. Participants completed four blocks of 66 trials each for a total of 264 trials.

An additional within-subject factor was tested in Experiment 3, object position, making it a 3 (scene-with-object/isolated object/scene-without-object) x 2 (original/swapped position) design. Each participant completed two blocks of 192 trials each for a total of 384 trials.

Within a block, each stimulus was repeated twice. The scene-with-object and isolated object condition were shown once with the object at its original position and once with the object at a swapped position. To avoid familiarity and carry-over effects, like previously, participants were presented with either the isolated object and scene-without-object version of a scene, or with the scene-with-object version of another scene. These two sets were counterbalanced across participants. Within each block the trial order was randomized.

#### EXPERIMENT 4

Before beginning the task, participants received a short demonstration of TMS, and the individual phosphene threshold was determined. The task procedure was identical to Experiments 1-3 except that after response (or timeout of 2 seconds), there was a variable inter-trial interval of 2 to 5 seconds (**Figure 2C**). This relatively long interval was chosen to prevent the coil from overheating and to prevent cumulative TMS effects (e.g., repetitive TMS); such an interval is commonly adopted in online TMS experiments (Gandolfo and Downing, 2019; Pitcher et al., 2008; 2007; Gandolfo et al., 2024). During the task, participants received two TMS pulses at 25Hz starting 60ms (early), 160ms (middle), or 260ms (late) relative to image onset. The stimulation onset was randomized across trials within each block and was not mentioned explicitly to the participants. Each TMS block (see Design below) lasted ∼3 minutes, with short breaks in between of ∼1 minute. The total duration of the experiment, including preparation and phosphene threshold determination, was approximately 90 minutes.

We used a 2×2×3 mixed design. The between-subjects factor was stimulation site (LOC, OPA) and the within-subjects factors included object presence (scene-with-object, scene-without-object) and stimulation onset (early, middle, late). Two TMS pulses could be applied randomly at one of three time-windows in every trial within a block: at 60ms and 100ms (early), 160ms and 200ms (middle), or at 260ms and 300ms (late) after stimulus onset. Every stimulus was repeated twice for each stimulation onset, resulting in 384 trials. The task was divided into 12 balanced blocks of 32 trials presented in random order.

### Analyses

#### EXPERIMENT 1-2

A repeated-measures ANOVA using a Greenhouse-Geisser sphericity correction was conducted to assess the effect of object presence on scene recognition accuracy (proportion of correct responses). Post-hoc pairwise comparisons were computed using a Bonferroni Correction, unless otherwise indicated. Analyses on reaction time and LISAS are reported in the **supplementary material**. If a one-sided binomial test comparing accuracy with 50% was insignificant (at α=.05), participants were considered at chance level and removed from the analyses.

#### EXPERIMENT 3

A 3×2 repeated measures ANOVA was conducted, with accuracy as the dependent variable and stimulus condition and object position as the independent variables. Since in the isolated scene condition the object position was dummy coded, we followed-up the interaction testing directly the 2 (original/swapped) x 2 (scene-with-object/isolated object) repeated measures ANOVA. Participants performing at chance level (based on a binomial test) were removed from the analyses.

#### EXPERIMENT 4

A 2×2×3 repeated measures ANOVAs was conducted on accuracy with stimulus condition (scene-with-object, scene-without-object), stimulation site (LOC, OPA), and stimulation onset (early, middle, late) as factors. Outliers were removed with a pre-registered criterion of being 2.5 standard deviations slower or less accurate than the overall mean across conditions. According to this criterion, four outliers were removed from the analysis and replaced to match the preregistered number of participants.

In accordance with our pre-registration (https://aspredicted.org/m9gp-k5hj.pdf), we tested the following planned comparisons contrasting performance for each onset and stimulation site: we expected LOC stimulation to decrease performance in the scene-with-object condition during stimulation in the middle timepoint (starting at 160ms) after stimulus presentation relative to the early time point (starting at 60ms). These effects on the LOC were expected to be specific for the scene-with-object condition and not observed in the scene-without-object condition. Following OPA stimulation, we expected effects on performance in both conditions, in the middle and late timepoint (starting from 160 ms after image onset) relative to the early timepoint. Due to the interplay between object and scene processing in the scene-with-object condition, we expected OPA stimulation effects to persist until the late stimulation timepoint. These hypotheses follow the logic and were pre-registered based on a previous chronometric TMS study on contextual *object* recognition (Wischnewski & Peelen, 2021A).

## RESULTS

### Experiment 1 - Objects facilitate scene recognition

We tested whether the presence of an object (originally belonging to the photograph) facilitates the recognition of otherwise hardly recognizable scenes. Participants performed indoor/outdoor judgments on briefly presented (50ms) photographs of 1) scenes with an object, 2) scenes without an object, or 3) an object alone on a uniform grey background (**Figure 2B**). Analyses on categorization accuracy showed a significant main effect of condition (*F*(1.857, 90.997) = 54.008, *p* <.001, partial η^2^ =.524) (**Figure 3A**). Scene classification accuracy was highest in the scene-with-object condition (*M* = 0.718, *SD* = 0.084) compared to both the scene-without-object (*M* = 0.644, *SD* = 0.082, *t*(49) *=* 6.818, *p_adj_* <.001, d = 0.95) and the isolated object conditions (*M* = 0.605, *SD* = 0.060, *t*(49) *=* 11.616, *p_adj_* <.001, d = 1.45). The scene and object alone conditions also differed, with the isolated-object condition showing the lowest performance (*t(49) =* 3.130, *p_adj_* =.009, d = 0.50).

**Figure 3.**
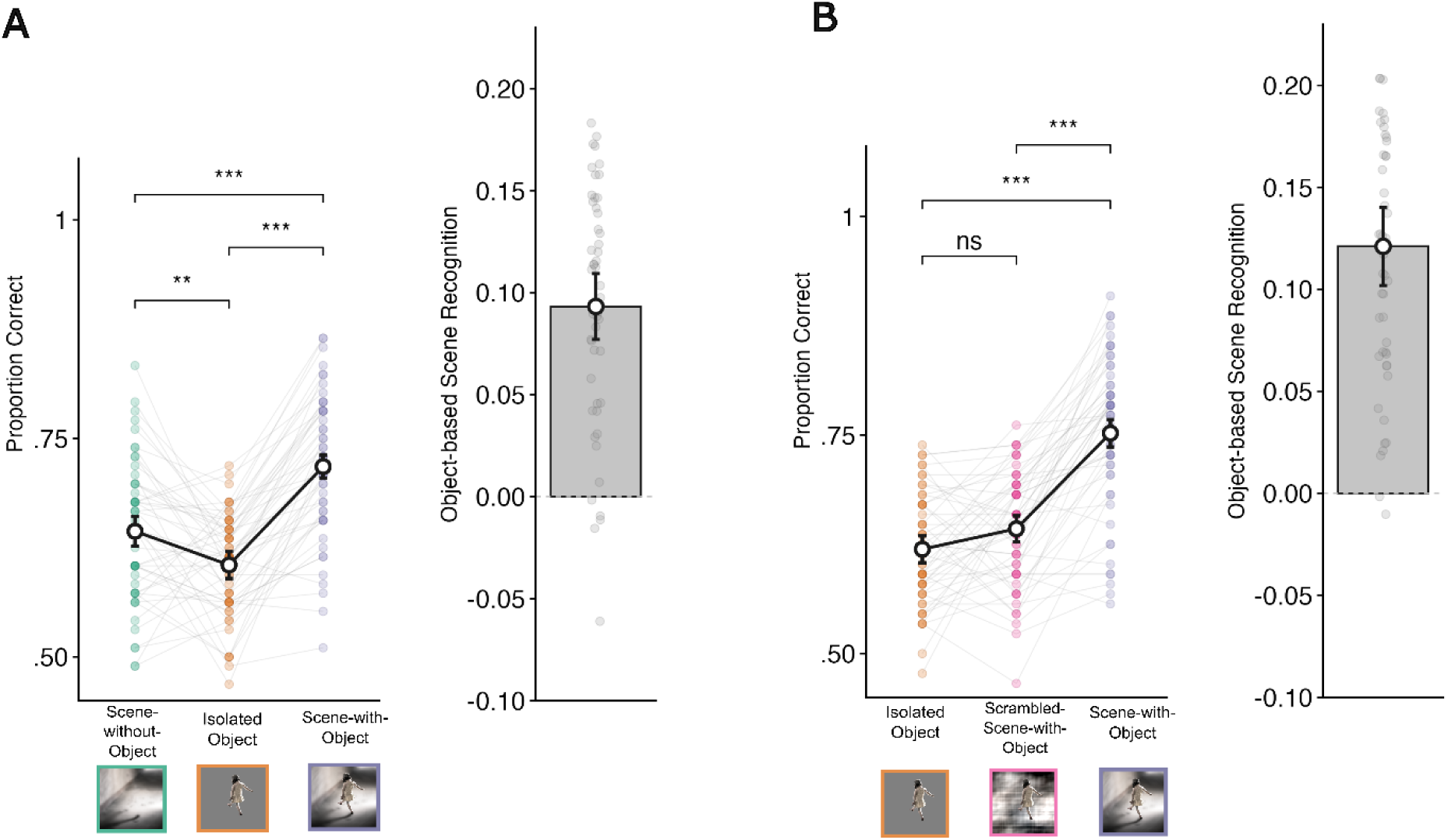
(A) Mean accuracy (expressed as proportion of correct responses) for Experiment 1 in each condition. Scene recognition accuracy was highest in the scene-with-object condition relative to the isolated and scene-without object conditions. (B) Mean accuracy for Experiment 2 in each condition. In both panels, the object-based scene recognition effect on the right is calculated as the difference between the scene-with-object condition and the mean of the remaining two conditions. Error bars represent within-subject 95% confidence intervals (Morey, 2008), points represent individual participants’ means. ns = not significant; \**p* ≤.05; \*\**p* ≤.01; \*\*\**p* ≤.001

**Figure 4.**
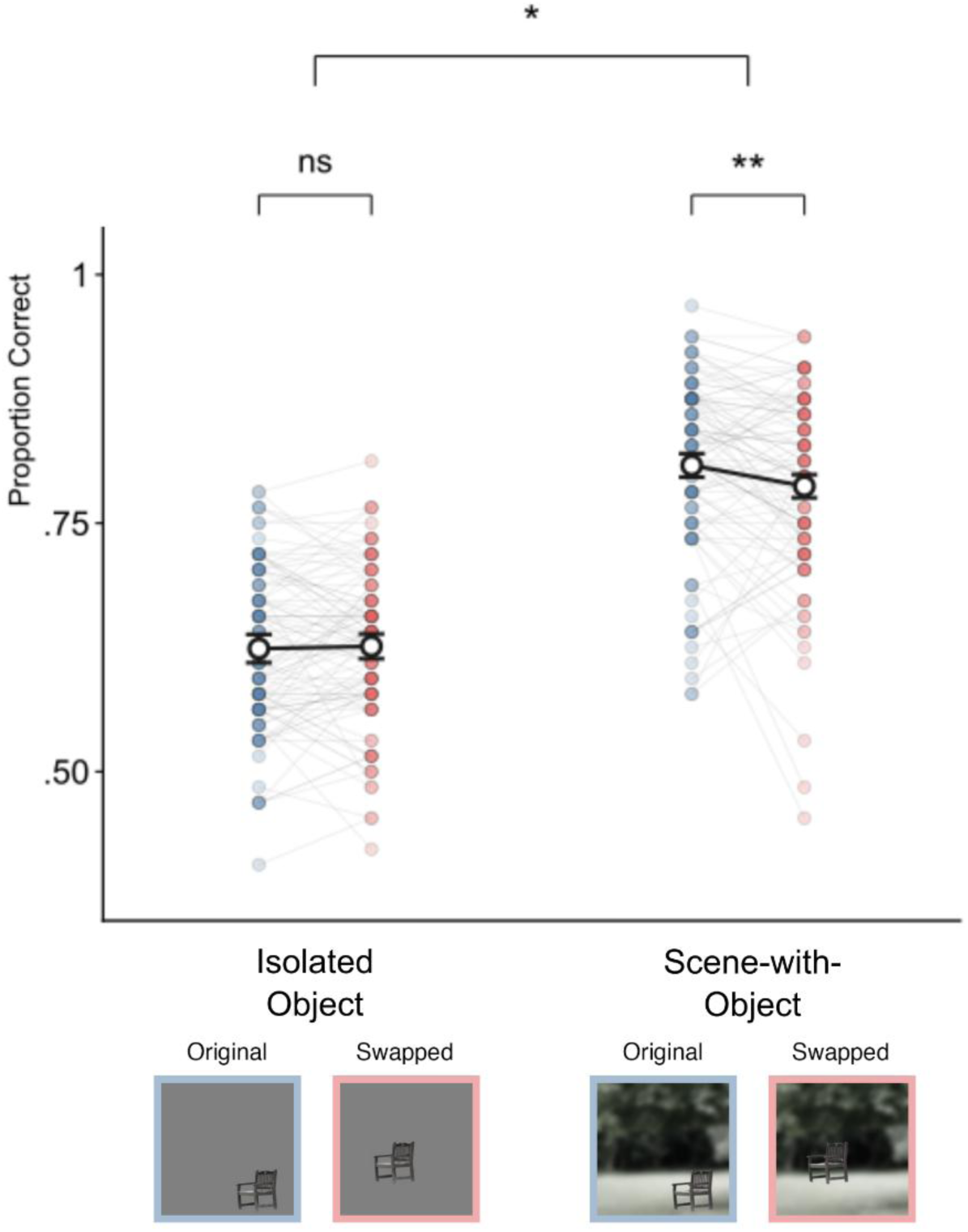
(A) Mean object recognition accuracy in each of the conditions. Scene recognition accuracy overall was higher in the scene-with-object condition than in the isolated object condition. In the scene-with-object condition, accuracy was higher when the object was at its original position. (B) The object-based scene recognition effect computed as the difference in accuracy between the scene-with-object condition and the isolated object condition separately for original and swapped object position. Error bars represent within-subject 95% confidence intervals (Morey, 2008), points represent individual participants’ means. \*\*\**p* ≤.001; \*\**p* ≤.01; \**p* ≤.05;

These results show that the presence of an object in a degraded scene facilitated scene recognition, showing that participants relied on object cues to achieve accurate scene categorization when the scene itself was not highly informative.

### Experiment 2 - Object-based scene recognition requires scene structure

In this experiment we sought to replicate the object-based scene recognition effect, this time contrasting categorization performance of the object in the degraded scene with the object being presented in a phase-scrambled version of that scene. Phase-scrambling preserves the overall contrast, colour, and luminance of a scene but disrupts scene structure. This is to rule out that the object-based recognition effect could arise via the mere co-occurrence between the objects and a scene’s low-level visual features. Scene recognition performance showed a main effect of condition (*F*(1.999, 99.777) = 81.277, p <.001 partial η^2^ =.619). Performance was best in the scene-with-object condition (*M* = 0.752, *SD* = 0.091) compared to both the scrambled-scene-with-object condition (*M* = 0.643, *SD* = 0.066, *t*(50) = 9.895, *p* <.001, d = 1.38) and the isolated object condition (*M* = 0.619, *SD* = 0.064, *t*(50) = 11.679, *p* <.001, d = 1.67). Accuracy did not differ between the isolated object and the scrambled-scene-with-object condition (*t*(50) = 2.141, *p* =.112, d = 0.30) (**Figure 3B**), indicating that removing scene structure through phase scrambling inhibits object-based scene recognition as much as an absent background.

### Experiment 3 - Object-based scene recognition relies on object position

In the previous experiments, participants performed above chance when the object was presented in isolation on a grey background (*M_E1_* = 0.605, *SD_E1_* = 0.060, *t_E1_*(49) = 13.349, *p_E1_* <.001; *M_E2_* = 0.619, *SD_E2_* = 0.064, *t_E2_*(50) = 13.349, *p_E2_* <.001). Despite the same object categories were used in both indoor and outdoor scenes, and that the object exemplar were selected to not be semantically informative of the scene, they likely still conveyed some information relevant for indoor/outdoor scene categorization (e.g., different types of chairs for indoor and outdoor environments). It is therefore possible that the object-based recognition effect of the first two experiments partly originated via the semantic information provided by the object (e.g., a person with a winter jacket on a white background is likely to be outdoor), rather than from a genuine facilitation in reading-out the surrounding space around the object. Here, we compared scene categorization accuracy capitalizing on another cue previously related to contextual object recognition, the object’s position within a scene (Biederman et al., 1982; Gayet et al., 2022). Object position provides information about scene layout (e.g., the location and orientation of the ground plane). We repeated the same task, this time randomly swapping the positions of the objects within the stimulus set (**Figure 1A**). We found an interaction effect between stimulus condition and object position on categorization accuracy (*F*(1,99) = 6.097, *p* =.029, partial η^2^ = 0.036). When the object was presented in the scene, performance was better with the object in its original position (*M* = 0.808, *SD* = 0.088) than at a swapped position (*M* = 0.787, *SD* = 0.094, *t*(99) = 2.798, *p* =.006, d = 0.25). The same position effect was not observed when the object appeared without the background (Original Position: *M* = 0.624, *SD* = 0.080; Swapped Position: *M* = 0.626, *SD* = 0.078, *t*(99) = 0.401, *p* =.689, d =.02) (**Figure 3A**). In **Figure 3B** we computed the object-based scene recognition index (subtracting the isolated object performance) separately for the original and swapped object positions.

These results show that the semantic contribution added by the object to the scene is not the only driver of object-based scene recognition. Rather, object appearance cues such as their original position in the scene further facilitate recognition of the scene.

### Experiment 4 - Causal contribution of object and scene selective regions in object-based scene recognition

The above experiments demonstrate that objects can facilitate the readout of scene cues for efficient scene recognition. Yet it remains unclear whether and when object-and scene-selective brain regions support the recognition of the scenes via the object. To address this question, we ran a pre-registered TMS experiment using chronometric online TMS. Using the same scene categorization task, we delivered TMS either to scene-selective OPA or object-selective LOC (in different participants) at three timepoints relative to scene onset (early – 60-100ms, middle - 160-200ms, late – 260-300ms), while participants were tasked to categorize either intact scenes or degraded scenes with their object. Importantly, we piloted the two types of scenes such that they would be comparable in terms of categorization performance at baseline, without stimulation (see Materials and Methods).

Our hypotheses were based on a recent TMS study investigating scene-based *object* recognition (Wischnewski and Peelen, 2021A). Briefly, in that study, when objects were categorized via a background scene, TMS to both object-selective LOC and scene-selective OPA reduced object categorization performance. Conversely, when isolated objects were categorized, LOC stimulation reduced performance, while OPA stimulation did not. These effects depended on stimulation onset: relative to an earlier stimulation onset (60-100ms, used as baseline), TMS to OPA impaired scene-based object recognition ∼160-200ms after image onset, while TMS to LOC impaired object recognition performance both at ∼160-200ms and later on (∼260-300ms), likely in virtue of feedback from scene processing. For recognition of isolated objects, the effects were observed on the LOC only, consistently with the time-course of object processing (160-200ms after image onset).

In the current study, we expected complementary stimulation and onset effects for object-based *scene* recognition, with recognition of scenes via the object requiring object-selective cortex. Indeed, we found a significant Stimulation Site x Object Presence interaction (*F*(1,47) = 6.071, *p* =.017, partial η^2^ =.114, **Figure 5AB**). Averaged across stimulation onset, TMS to LOC reduced scene categorization performance in the scene-with-object condition relative to the recognition of isolated scenes, (*t*(23) =-2.561, *p* =.014, d = 0.484 - **Figure 5A**). By contrast, OPA stimulation did not differentially reduce scene categorization in the scene-with-object and the scene-without-object conditions (*t*(24) = 0.907*, p* =.369, d = 0.168). n.

**Figure 5.**
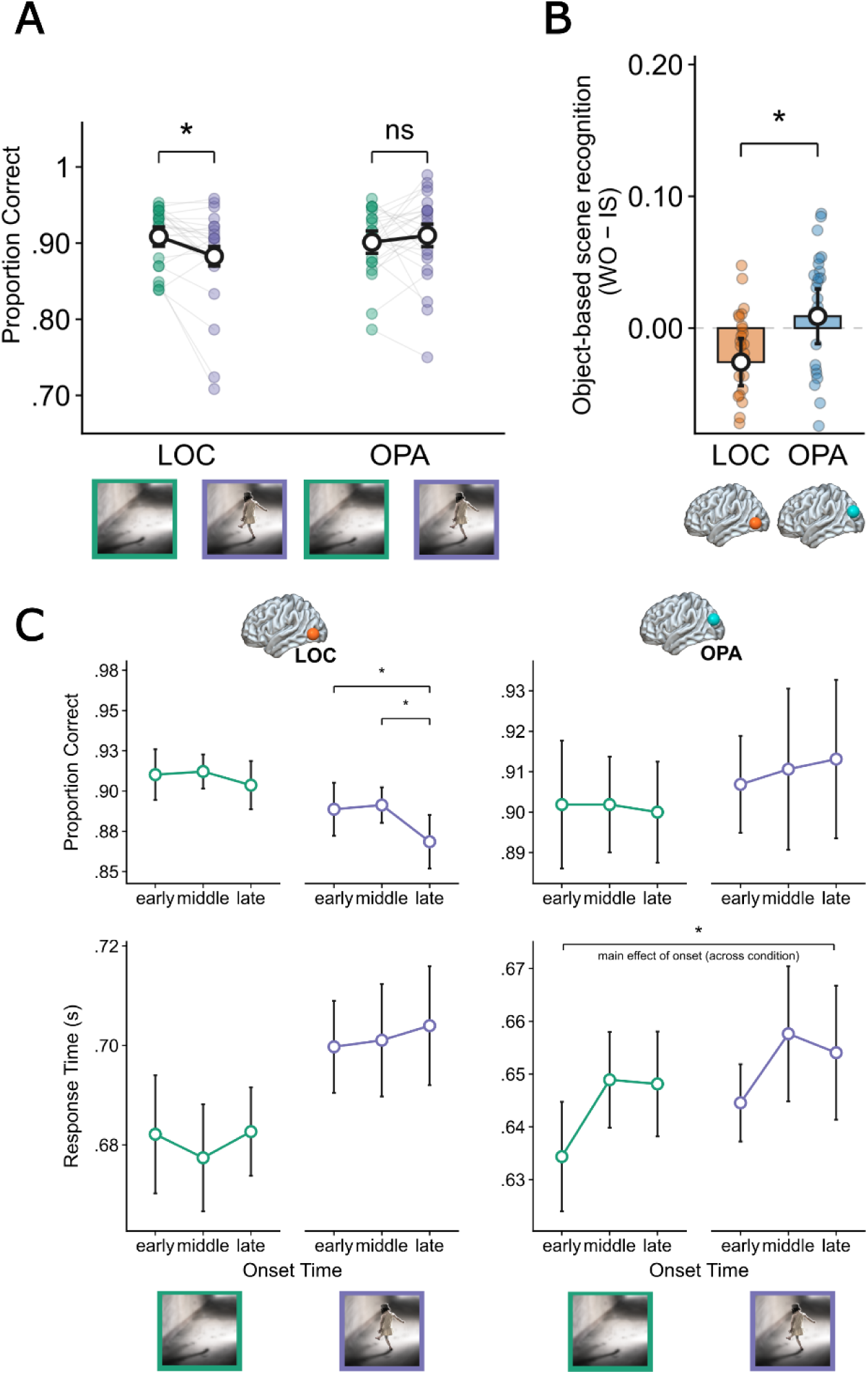
(A) Average performance across stimulation onset for each stimulation site in the scene-with-object and scene-without-object conditions. Only after LOC stimulation, accuracy decreased in the scene-with-object condition compared to the scene-without-object condition. During OPA stimulation, no difference between stimulus conditions occurred. (**B**) Object-based scene recognition calculated separately for each stimulation site expressing the site x condition interaction. Object presence decreased performance during LOC stimulation relative to OPA stimulation. \*\*\**p* ≤.001; \*\**p* ≤.01; \**p* ≤.05; ns = not significant. (**C**) Average performance in accuracy (upper quadrants) and response times (lower quadrants). In accordance with the pre-registration, we tested: 1) onset effects of the LOC in the scene-with-object condition (in purple). Here, performance decreased in terms of accuracy in the late time window relative to the earlier time windows. 2) onset effects across condition during OPA stimulation. Here, performance was affected via slowing down responses in the middle and late time windows relative to the early one across conditions. Error bars represent within-subject 95% confidence intervals (Morey, 2008), points represent individual participants’ means.

We expected but did not find reliable onset effects (no three-way interaction of Stimulation Site x Stimulation Onset x Object Presence (*F*(2,94) = 0.866, *p* =.424, partial η^2^ =.018). Following our pre-registered analyses, we tested if there was an onset effect on the LOC specifically in the scene-with-object condition. There was a main effect of Stimulation Onset (*F*(1.906, 43.840) = 3.794, *p* = 0.032, partial η^2^ = 0.142). Accuracy was lower in the late time-window relative to the early (*t*(23) = 2.50, *p* = 0.020, d = 0.51) and middle (*t*(23) = 2.47, *p* = 0.021, d = 0.50) time-windows and did not reliably differ between the early and middle time window (*t*(23) = 0.27, *p* = 0.793, d = 0.05). The same main effect of onset did not reach significance for the reaction times (*F*(1.951, 44.873) = 0.203, *p =* 0.811, partial η^2^ = 0.009). In the scene-without-object condition, stimulation of LOC did not affect performance at any specific time-window, neither in accuracy (main effect of onset: *F*(1.951, 44.872) = 0.619, *p* = 0.539, partial η^2^ = 0.026) nor reaction times (main effect of onset: *F*(1.943, 44.692) = 0.398, *p* = 0.668, partial η^2^ = 0.017). These results show that LOC stimulation effects in the scene-with-object condition were particularly evident in the late time window after image onset relative to the earlier ones (**Figure 5C**).

Based on our pre-registration, we expected stimulation of OPA to affect scene judgments in both conditions, with onset effects sparing the early time window, and to persist in the late time window (in virtue of feedback from the LOC) specifically in the scene-with-object condition. Across the two conditions, we did not observe a main effect of onset on the categorization accuracy (*F*(1.606, 38.545) = 0.079, *p =* 0.886, partial η^2^ =.003). A main effect of Stimulation Onset was observed on response times (*F*(1.762, 42.280) = 4.921, *p* = 0.015, partial η^2^ =.170). A simple effect analysis showed that relative to the early time-point participants were slower at categorizing scenes in the middle (*t*(24) = 2.831, *p* = 0.009, d = 0.363), and late timepoint (*t*(24) = 2.160, *p* = 0.040, d = 0.306). For both dependent variables, there was no significant stimulation by onset interaction (accuracy: *F*(1.989, 47.724) = 0.211, *p* = 0.809, partial η^2^ =.009; response times: *F*(1.689, 40.540) = 0.195, *p* = 0.787, partial η^2^ =.008) meaning that the onset effects of stimulation similarly affected both scene conditions regardless of their object, and in both cases started 160ms after image onset (**Figure 5C**).

### General Discussion (max 1500)

In this study, we provide evidence for object-based scene recognition: scene categorization improved when an object was placed in an otherwise hardly recognizable scene. This benefit depended on the availability of the scene’s structural cues, rather than on its low-level properties (colour, overall contrast, and luminance), and on the object’s position within the scene, beyond its semantic informativeness. Finally, we report a causal involvement of the LOC in object-based scene recognition. Together, these findings show how, by disambiguating scene layout, objects can facilitate scene recognition.

Objects placed in ambiguous scenes improved indoor/outdoor scene categorization, an effect suggesting an object-based scene disambiguation process. By disambiguation, we mean the process through which contextual information helps resolve the identity of a stimulus that is itself poorly visible or ambiguous. In the scene-object interactions literature, this contextual facilitation is thought to operate at a perceptual stage: when a stimulus is degraded, the context enhances its representation in visual cortex, effectively sharpening the percept (for a review, see Peelen et al., 2024). The mechanism has been most clearly demonstrated for scene-based object recognition, where scene context boosts the cortical representation of degraded objects (Brandman & Peelen, 2017; Leticevscaia et al., 2024), but it has also been reported in the reverse direction, with diagnostic objects enhancing the representation of ambiguous scene layouts in scene-selective cortex (Brandman & Peelen, 2019, 2023).

While previous research on object-based scene recognition used objects that are strongly indicative of scene category within intact scenes (Davenport & Potter, 2004; Davenport, 2007), we show that objects can disambiguate the surrounding layout even when they are not strongly semantically congruent with their background. This suggests that, when a scene itself contains little information, scene classification can be achieved through object processing. The object-based scene recognition effect did not occur when the object appeared in a phase-scrambled scene (Experiment 2), indicating that co-occurrence between the low-level properties of the scene and the object is not sufficient to facilitate scene judgments. Rather, the disambiguating object supports a better encoding of the properties that define a scene’s structure.

In the first two experiments, however, performance for objects shown without scenes remained above chance. As such, the object benefit for scene judgments could have arisen from a summation of the semantic information carried by the object and the scene when presented together, rather than from genuine disambiguation of the scene by the object. Experiment 3 ruled this out: the mere co-occurrence of object and scene did not fully account for the object-based scene recognition effect, which was reduced when the object’s original location was shifted within the photograph. Compared with previous studies of object-based scene recognition, we thus demonstrate that an object’s position in the scene contributes to scene recognition in its own right, independently of its identity. A cat, for example, is equally likely to be seen outdoors or indoors (see **Figure 1**). Yet, the way in which it appears, how it is sitting, its size and the position in which it is standing, further define the surrounding scene, adding depth information and perspective that results in improved scene recognition.

Together, these findings show that object-based disambiguation for scene recognition shares properties with the context effects typically studied in object recognition. Scene disambiguation via the object relied on both semantic and syntactic associations between objects and scenes (Biederman et al., 1982; Võ & Wolfe, 2013; Kaiser et al., 2019; Võ et al., 2019; Võ, 2021). An open question for future research is the extent to which scene disambiguation depends on how diagnostic an object is, both semantically and syntactically, for recognizing a given scene, and whether the same object cues can support different types of scene categorization (for example, recognizing a place not merely as indoor or outdoor, but as a kitchen or a bathroom).

### Causal evidence for object-based scene disambiguation in visual cortex

Experiment 4 showed that efficient object-based scene recognition relies on the object-selective LOC. TMS over the left LOC selectively decreased performance in the scene-with-object condition while leaving the isolated-scene condition unaffected. This is consistent with fMRI work suggesting a supporting role of the object-selective LOC in scene recognition (MacEvoy & Epstein, 2011) and provides the first causal evidence that the LOC contributes to efficient scene judgments. When the ambiguity of the scene pushes observers to recruit object-related information to solve a scene task, object-specific visual processing becomes causally necessary.

Contrary to our predictions, however, the effects of stimulation onset were less clear, as they were not supported by an interaction between stimulation site, onset, and condition. We had expected a clear chronometry, whereby disambiguation of the scene via the object would require object processing around 160 ms after image onset (Wischnewski & Peelen, 2021A). Instead, the preregistered contrasts revealed that LOC stimulation reduced accuracy in the scene-with-object condition particularly in the latest time window (260 ms after onset). It is possible that the greater task-relevance of scene over object recognition, and/or the position of the objects away from fixation, delayed the involvement of scene processing in the task.

The stimulation effects disrupting scene-selective processing were observed in response times: OPA stimulation delayed scene recognition in the middle and late time windows, relative to the early window, across all conditions. Scene processing thus began before, and persisted throughout, object processing. We had expected that, as a consequence of object disambiguation, scene processing would become further engaged in the scene-with-object condition at a later time window, once the LOC had completed feedforward object processing. However, because the LOC effects peaked in a late time window, any scene-related activity following object processing may fall outside the onsets we tested. As such, the present data cannot establish whether the information disambiguated by the object was fed back to scene-selective cortex. Consistent with this possibility, a recent MEG study found that object-based scene disambiguation peaks around 320 ms and persists until about 350 ms after stimulus onset (Brandman & Peelen, 2023). It is worth noting that these onset effects were reported in keeping with the pre-registration but did not survive corrections for multiple comparisons. As such, further stimulation studies may better characterize the chronometry of object-based scene recognition.

Nonetheless, Experiment 4 clearly shows that, overall, LOC stimulation reduces accuracy in object-based scene recognition, even after performance has been equated with the scene-without-object condition. It remains an open question which components of object processing the stimulation disrupted (for example, the semantic component, the object’s appearance, or both). Future studies could test whether LOC stimulation specifically affects the scene-with-object condition when it is contrasted with a phase-scrambled scene containing the object, with the position effect, or with other object features relevant to scene disambiguation (such as retinal size; Gayet et al., 2022). Including a later time window could likewise clarify whether scene processing continues after the disambiguating object has been processed, mirroring what has been reported for scene-based object recognition (Wischnewski and Peelen, 2021A).

## Conclusions

In this study, we show that when people view an ambiguous scene, they use object-specific information to disambiguate its layout in order to judge whether it is indoor or outdoor. This effect is not fully explained by semantic associations between object and scene; it also reflects the physical properties of the object (such as its position) and of the scene (such as its structure). Finally, using transcranial magnetic stimulation, we show that object-selective visual cortex is causally involved in scene recognition when object information is used to disambiguate the scene’s layout. Together, these findings demonstrate that an object’s visual properties, including its position, size, and orientation, can disambiguate the surrounding scene to support scene categorization.

## Author Contribution

Aaron Schnippe: Conceptualization, Methodology, Formal Analysis, Data Curation, Investigation, Resources, Software, Visualization, Writing-Original Draft, Writing – Review and Editing

Natalia Rutkowska: Conceptualization, Investigation, Resources, Visualization, Writing – Review and Editing, Methodology

Marius V. Peelen: Conceptualization, Methodology, Funding acquisition, Supervision, Writing – Review and Editing

Marco Gandolfo: Conceptualization, Methodology, Formal Analysis, Validation, Resources, Software, Funding acquisition, Supervision, Software, Project Administration, Writing – Original Draft, Writing – Review and Editing

## Acknowledgments

We thank the PeelenLab members for useful feedback on the project and the Technical Support Group of the Faculty of Social Sciences at Radboud University for technical advice. We also thank Miles Wischnewski and Surya Gayet for Matlab code resources.

## Funding information

This project has received funding from the European Union’s Horizon 2020 research and innovation program under the Marie Skłodowska-Curie fellowship (grant agreement No. 101033489) and from the European Research Council (ERC) under the European Union’s Horizon 2020 research and innovation program (Grant Agreement No. 725970).

## Open Practices statement

The raw data and stimuli generated from this project are publicly available at the Open Science Framework repository – https://osf.io/teuxv/overview?view_only=d46fb6f0a8784dcf9e19c6880bf228c9

## Supplementary Materials

### Analyses of reaction time and LISAS

In the reaction time analysis, only reaction times of correct trials were considered. The linear integrated speed-accuracy score is defined as:

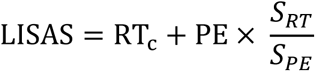

Where RT_C_ is the average reaction time in correct trials, PE is the proportion of errors (defined as 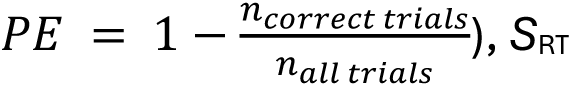 refers to the standard deviation of the correct RTs and *S*_PE_ refers to the standard deviation of the proportion of errors (Vandierendonck, 2017; 2018). Lower LISAS indicates better performance.

### Experiment 1

The reaction times in accurate trials showed no effect of condition (*F*(1.842, 90.350) = 0.609, *p* =.533, partial η^2^ =.012). However, when considering performance as a combination of speed and accuracy, i.e. the LISAS, the effect of condition was significant (*F*(1.938, 94.949) = 10.939, *p* <.001, partial η^2^ =.182). Post-hoc pairwise comparisons confirmed a similar pattern to the accuracy results, ruling out that the pattern observed would be explained by a speed-accuracy trade-off. Indeed, LISAS showed a better performance in the scene-with-object condition (*μ* = 0.874, *SD* = 0.293) than in the isolated object condition (*μ* = 0.925, *SD* = 0.282, *t*(49) = 4.891, *p_adj_* <.001, d = 0.64), and better performance in the scene-with-object condition than in the isolated scene condition (*μ* = 0.912, *SD* = 0.306, *t*(49) = 3.154, *p_adj_* =.008, d = 0.49). LISAS did not differ between the isolated scene and isolated object condition (*t*(49) = 1.099, *p_adj_* =.831, d = 0.16). LISAS was lowest for the scene-with-object condition, indicating best performance.

### Experiment 2

Reaction time did not differ based on condition (*F*(1.989, 99.444) = 2.547, *p* =.084, partial η^2^ =.048). There was a significant effect of condition on LISAS (*F*(1.941, 95.097) = 25.825, *p* <.001, partial η^2^ =.345). Post-hoc pairwise comparisons with a Bonferroni correction revealed that there were significant differences in LISAS between the scene-with-object (*μ* = 0.773, *SD* = 0.170) and scrambled-scene-with-object condition (*μ* = 0.814, *SD* = 0.161, *t*(49) = 5.399, *p_adj_* >.001, d = 0.77) and between the scene-with-object and isolated object condition (*μ* = 0.825, *SD* = 0.176, *t*(49) = 6.346, *p_adj_* >.001, d = 0.96), but not between the isolated object and scrambled-scene-with-object condition (*t*(49) = 1.492, *p_adj_* =.426, d = 0.19), supporting the results of the previous analysis on accuracy.

### Experiment 3

To precisely assess the effect of object position, the isolated scene condition was removed from the following analyses since its stimuli did not include an object. In the 2×2 repeated measures ANOVA predicting reaction time from stimulus condition and object position, a main effect of condition was found (*F*(1,99) = 36.084, *p* <.001, partial η^2^ = 0.267). A simple main effect analysis showed that participants responded quicker in the scene-with-object condition (*μ* = 0.656, *SD* = 0.084) than in the isolated object condition (*μ* = 0.686, *SD* = 0.109, *t* = 6.007, *p* <.001). The effect size, as measured by Cohen’s d, was d =.62. An interaction effect of position and condition on LISAS was found (*F*(1,99) = 5.133, *p* =.026, partial η^2^ = 0.049). A simple interaction effect analysis revealed that LISAS differed between positions within the scene-with-object condition (μ_original_ = 0.730, *SD*_original_ = 0.110, μ_swapped_ = 0.742, *SD*_swapped_= 0.122, *t*(99) = 2.498, *p* =.014, d =.16), but not in the isolated object condition (μ_original_ = 0.828, *SD*_original_ = 0.152, μ_swapped_= 0.827, *SD*_swapped_= 0.160, t(99) = 0.273, *p* =.785, d =.01). In the scene-with-object condition, LISAS was lower for the original object position, indicating better performance when the object was at its original position as compared to a random other position in the scene. These findings support the results of the accuracy analysis and show that that effect cannot be explained by a speed-accuracy trade-off.

### Experiment 4

In the reaction times analysis, we found a significant main effect of stimulus condition (*F*(1,47) = 9.213, *p* =.004, partial η^2^ =.164). Reaction times were quicker in the scene-without-object condition (*M* = 0.662, *SD* = 0.101) than in the scene-with-object condition (*M* = 0.677, *SD* = 0.106, *t*(48) = 3.494, *p* =.001, d = 0.39). We found no significant main effect of stimulation onset (*F*(2,94) = 2.155, *p* =.122, partial η^2^ =.044) nor a three-way interaction between stimulation site, stimulus onset and object presence (*F*(2,94) = 0.406, *p* =.667, partial η^2^ =.009).

On LISAS, we found a significant two-way interaction between stimulation site and stimulus condition (*F*(1,47)=4.395, *p* =.041, partial η^2^ =.086), ruling out speed accuracy trade-offs on the main analysis. A simple effects analysis showed that, during LOC stimulation, participants had lower LISAS (i.e., better performance) in the scene-without-object condition than in the scene-with-object condition (*t*(47) = 3.048, *p* =.004, d = 0.23). This difference in performance between conditions was not found during OPA stimulation (*t*(47) = 0.116, *p* =.908, d = 0.01). Comparing performance within stimulus conditions between stimulation sites, we find that performance in the scene-without-object condition does not differ depending on stimulation site (*t*(47) = 1.512, *p* =.137, d = 0.40). However, in the scene-with-object condition, performance is better during OPA stimulation than during LOC stimulation (*t*(47) = 2.197, *p* =.033, d = 0.62). This finding further supports the previous findings that the LOC is involved in the recognition of the scene, and crucially, that this recognition is achieved via object processing.

### Pre-registered planned comparisons on LISAS

Following our pre-registered analyses, we tested for effects of stimulation onset on LISAS during LOC stimulation within the scene-with-object condition. We found no main effect of stimulation onset (*F*(1.768, 40.656) = 1.955, *p* = 0.159, partial η^2^ = 0.078). Our planned pairwise comparisons revealed no differences in LISAS between early and middle (*t*(23) = 0.487, *p* =.631, d = 0.11), middle and late (*t*(23) = 1.658, *p* =.111, d = 0.28) or early and late stimulation onset (*t*(23) = 1.906, *p* =.069, d = 0.39).

Further, we investigated stimulation onset effects on LISAS during OPA stimulation across object presence conditions. We found no main effect of stimulation onset (*F*(1.897, 45.529) = 1.740, *p* = 0.188, partial η^2^ = 0.068). Performance did not differ between early and middle (*t*(24) = 1.639, *p* =.114, d = 0.21), middle and late (*t*(24) = 0.310, *p* =.759, d = 0.04) or early and late stimulation onset (*t*(24) = 1.630, *p* =.116, d = 0.17).

### Stimulation site selection Procedure – Experiment 4

The stimulation-site selection procedure took place 3 to 7 days prior to the experiment. Participants received four-pulse TMS over LOC, OPA or vertex while performing a scene recognition (indoor/outdoor) and an object recognition task (7 object categories). The object recognition task was adapted from Wischnewski and Peelen (2021B). The individual data resulting from this session were used to assign participants stimulation site in the chronometric TMS study. For an overview of two TMS parts, see **Figure S1**. This selection procedure was performed to increase the chances of finding chronometric TMS effects (Wischnewski & Peelen 2021A, 2021B).

### Localization procedure: Transcranial Magnetic Stimulation protocol

The localization procedure used the same TMS set-up as the chronometric TMS study with the following changes. Four TMS pulses (biphasic, wavelength: 280 µs) at 10 Hz were delivered online at a fixed intensity of 60% of the maximum stimulator output. This four pulse TMS design was chosen to disrupt the functioning of the selected areas for the whole duration of the perception process. TMS was applied over left LOC, left OPA or vertex. The vertex acted as a control and was individually determined at half of the distance between inion and nasion.

---

**Figure S1.**
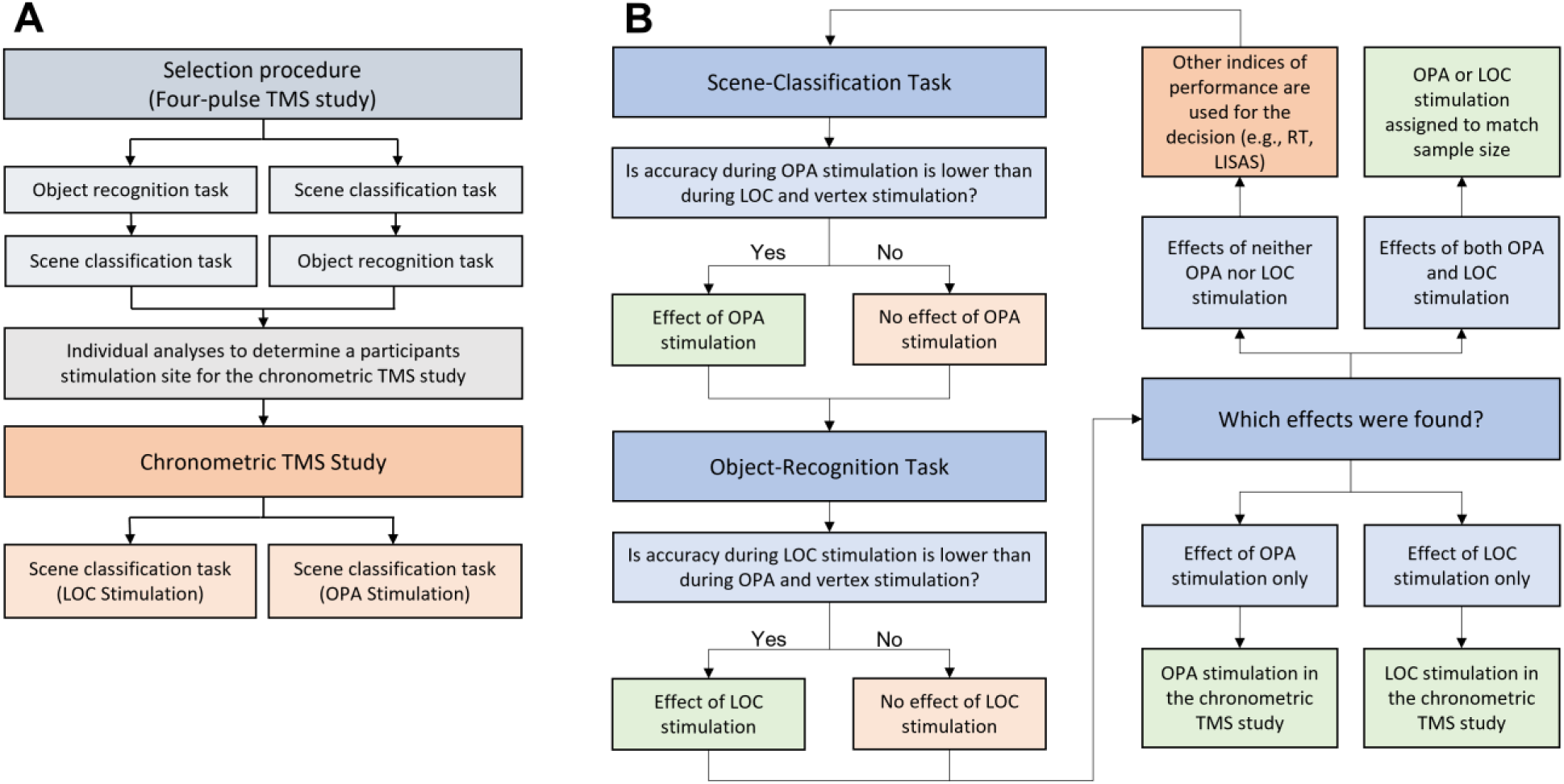
(A) Overview of the two TMS studies. In the site selection procedure, participants completed an object recognition task and a scene classification task (counterbalanced across participants). Individual analyses were conducted to assign participants to a stimulation site condition in the chronometric TMS study. The chronometric TMS study, took place three to seven days after the selection procedure. (B) Overview of the assignment of the stimulation size for the chronometric TMS study. To increase the chances of finding a chronometric TMS effects, participants were assigned the stimulation site in the chronometric TMS part depending on the results of this functional localization procedure. In the object recognition task, performance was first compared between LOC and OPA, and then between LOC and vertex. Participants that showed lower accuracy in the object recognition task during LOC stimulation when compared to OPA and vertex were then assigned to receive LOC stimulation in the chronometric TMS study. In the scene classification task, performance was first compared between OPA and LOC, and then between OPA and vertex. Participants that showed lower accuracy in the scene classification task during OPA stimulation when compared to LOC and vertex were then assigned to receive OPA stimulation in the chronometric TMS study. Participants showing both an effects of LOC stimulation in the object recognition task and an effect of OPA stimulation in the scene classification task were assigned to equate sample sizes across stimulation sites in the chronometric TMS study. If no effects of stimulation on accuracy was found, other indicators of performance, such as mean reaction time or LISAS, were used for assignment.

## Notes

### Competing Interest Statement

The authors have declared no competing interest.

