## supplemental_Materials for "Behavioral and causal evidence for object-based scene recognition in visual cortex"

### Supplementary Materials

#### Analyses of reaction time and LISAS

In the reaction time analysis, only reaction times of correct trials were considered. The linear integrated speed-accuracy score is defined as:

$$\mathrm{LISAS} = \mathrm{RT_c} + \mathrm{PE} \times \frac{S_{RT}}{S_{PE}}$$

Where $RT_C$ is the average reaction time in correct trials, PE is the proportion of errors (defined as $PE = 1 - \frac{n_{correct\ trials}}{n_{all\ trials}}$), $S_{RT}$ refers to the standard deviation of the correct RTs and $S_{PE}$ refers to the standard deviation of the proportion of errors (Vandierendonck, 2017; 2018). Lower LISAS indicates better performance.

#### Experiment 1

The reaction times in accurate trials showed no effect of condition ($F(1.842, 90.350) = 0.609$, $p = .533$, partial $\eta^2 = .012$). However, when considering performance as a combination of speed and accuracy, i.e. the LISAS, the effect of condition was significant ($F(1.938, 94.949) = 10.939$, $p < .001$, partial $\eta^2 = .182$). Post-hoc pairwise comparisons confirmed a similar pattern to the accuracy results, ruling out that the pattern observed would be explained by a speed-accuracy trade-off. Indeed, LISAS showed a better performance in the scene-with-object condition ($\mu = 0.874$, $SD = 0.293$) than in the isolated object condition ($\mu = 0.925$, $SD = 0.282$, $t(49) = 4.891$, $p_{adj} < .001$, d = 0.64), and better performance in the scene-with-object condition than in the isolated scene condition ($\mu = 0.912$, $SD = 0.306$, $t(49) = 3.154$, $p_{adj} = .008$, d = 0.49). LISAS did not differ between the isolated scene and isolated object condition ($t(49) = 1.099$, $p_{adj} = .831$, d = 0.16). LISAS was lowest for the scene-with-object condition, indicating best performance.

#### Experiment 2

Reaction time did not differ based on condition ($F(1.989, 99.444) = 2.547$, $p = .084$, partial $\eta^2 = .048$). There was a significant effect of condition on LISAS ($F(1.941, 95.097) = 25.825$, $p < .001$, partial $\eta^2 = .345$). Post-hoc pairwise comparisons with a Bonferroni correction revealed that there were significant differences in LISAS between the scene-with-object ($\mu = 0.773$, $SD = 0.170$) and scrambled-scene-with-object condition ($\mu = 0.814$, $SD = 0.161$, $t(49) = 5.399$, $p_{adj} > .001$, d = 0.77) and between the scene-with-object and isolated object condition ($\mu = 0.825$, $SD = 0.176$, $t(49) = 6.346$, $p_{adj} > .001$, d = 0.96), but not between the isolated object and scrambled-scene-with-object condition ($t(49) = 1.492$, $p_{adj} = .426$, d = 0.19), supporting the results of the previous analysis on accuracy.

#### Experiment 3

To precisely assess the effect of object position, the isolated scene condition was removed from the following analyses since its stimuli did not include an object. In the 2x2 repeated measures ANOVA predicting reaction time from stimulus condition and object position, a main effect of condition was found ($F(1,99) = 36.084$, $p < .001$, partial $\eta^2 = 0.267$). A simple main effect analysis showed that participants responded quicker in the scene-with-object condition ($\mu = 0.656$, $SD = 0.084$) than in the isolated object condition ($\mu = 0.686$, $SD = 0.109$, $t = 6.007$, $p < .001$). The effect size, as measured by Cohen's d, was d = .62. An interaction effect of position and condition on LISAS was found ($F(1,99) = 5.133$, $p = .026$, partial $\eta^2 = 0.049$). A simple interaction effect analysis revealed that LISAS differed between positions within the scene-with-object condition ($\mu_{original} = 0.730$, $SD_{original} = 0.110$, $\mu_{swapped} = 0.742$, $SD_{swapped} = 0.122$, $t(99) = 2.498$, $p = .014$, d = .16), but not in the isolated object condition ($\mu_{original} = 0.828$, $SD_{original} = 0.152$, $\mu_{swapped} = 0.827$, $SD_{swapped} = 0.160$, t(99) = 0.273, $p = .785$, d = .01). In the scene-with-object condition, LISAS was lower for the original object position, indicating better performance when the object was at its original position as compared to a random other position in the scene. These findings support the results of the accuracy analysis and show that that effect cannot be explained by a speed-accuracy trade-off.

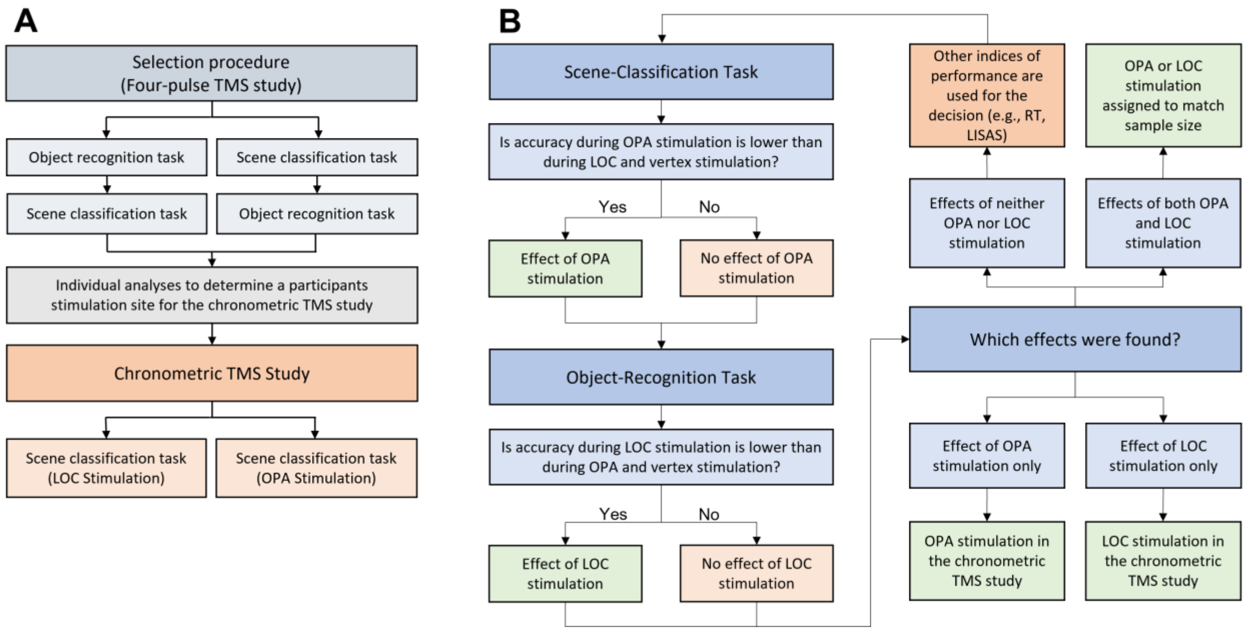


**Figure S1**. (A) Overview of the two TMS studies. In the site selection procedure, participants completed an object recognition task and a scene classification task (counterbalanced across participants). Individual analyses were conducted to assign participants to a stimulation site condition in the chronometric TMS study. The chronometric TMS study, took place three to seven days after the selection procedure. (B) Overview of the assignment of the stimulation size for the chronometric TMS study. To increase the chances of finding a chronometric TMS effects, participants were assigned the stimulation site in the chronometric TMS part depending on the results of this functional localization procedure. In the object recognition task, performance was first compared between LOC and OPA, and then between LOC and vertex. Participants that showed lower accuracy in the object recognition task during LOC stimulation when compared to OPA and vertex were then assigned to receive LOC stimulation in the chronometric TMS study. In the scene classification task, performance was first compared between OPA and LOC, and then between OPA and vertex. Participants that showed lower accuracy in the scene classification task during OPA stimulation when compared to LOC and vertex were then assigned to receive OPA stimulation in the chronometric TMS study. Participants showing both an effects of LOC stimulation in the object recognition task and an effect of OPA stimulation in the scene classification task were assigned to equate sample sizes across stimulation sites in the chronometric TMS study. If no effects of stimulation on accuracy was found, other indicators of performance, such as mean reaction time or LISAS, were used for assignment.

#### *Localization procedure: Transcranial Magnetic Stimulation protocol*

The localization procedure used the same TMS set-up as the chronometric TMS study with the following changes. Four TMS pulses (biphasic, wavelength: 280 μs) at 10 Hz were delivered online at a fixed intensity of 60% of the maximum stimulator output. This four pulse TMS design was chosen to disrupt the functioning of the selected areas for the whole duration of the perception process. TMS was applied over left LOC, left OPA or vertex. The vertex acted as a control and was individually determined at half of the distance between inion and nasion.
